# Structural basis of membrane recognition of *Toxoplasma gondii* vacuole by Irgb6

**DOI:** 10.1101/2021.07.01.450801

**Authors:** Yumiko Saijo-Hamano, Aalaa Alrahman Sherif, Ariel Pradipta, Miwa Sasai, Naoki Sakai, Yoshiaki Sakihama, Masahiro Yamamoto, Daron M Standley, Ryo Nitta

## Abstract

The p47 immunity-related GTPase (IRG) Irgb6 plays a pioneering role in host defense against *Toxoplasma gondii* infection. It is recruited to the parasitophorous vacuole membrane (PVM) formed by *T. gondii* and disrupts it. Despite the importance of this process, the molecular mechanisms accounting for PVM recognition by Irgb6 remain elusive due to lack of structural information on Irgb6. Here we report the crystal structures of mouse Irgb6 in the GTP-bound and nucleotide-free forms. Irgb6 exhibits a similar overall architecture to other IRGs in which GTP-binding induces conformational changes in both the dimerization interface and the membrane-binding interface. The membrane-binding interface of Irgb6 assumes a unique conformation, composed of N- and C-terminal helical regions forming a phospholipid binding site. *In silico* docking of phospholipids further revealed membrane binding residues that were validated through mutagenesis and cell-based assays. Collectively, these data demonstrate a novel structural basis for Irgb6 to recognize *T. gondii* PVM in a manner distinct from other IRGs.

**Summary:** Upon *Toxoplasma gondii* infection, Irgb6 is recruited to the parasitophorous vacuole membrane (PVM) where it disrupts it. We solved the atomic structures of Irgb6 in two distinct nucleotide states, revealing a unique PVM binding interface sensitive to the GTPase cycling.

## Introduction

Infection by intracellular pathogens stimulates innate and acquired immune systems to produce interferon (IFN). IFN-γ is a proinflammatory cytokine produced from natural killer cells and T cells. Binding of IFN-γ with IFN-γ receptors activates gene expression programs via the JAK-STAT pathways. A number of IFN-γ -inducible products play pivotal and pleiotropic roles in cell-autonomous immunity against various intracellular pathogens such as viruses, bacteria and protozoan parasites (MacMicking, 2012).

*Toxoplasma gondii* is an important human and animal pathogen that causes lethal toxoplamosis in immune-compromised individuals such as those receiving bone marrow transplantations or suffering from AIDS (Boothroyd, 2009; Goldstein et al., 2008). IFN-γ suppresses intracellular *T. gondii* growth in a manner dependent on inducible nitric oxide production by nitric oxide synthase 2 (NOS2) and tryptophan degradation by indoleamine 2,3-deoxygenase (IDO), both of which are important for prevention of chronic toxoplasmosis (Divanovic et al., 2012; Sasai et al., 2018; Scharton-Kersten et al., 1997). In contrast, recent studies demonstrate that host defense during acute toxoplamosis requires IFN-γ -inducible GTPases that localize at a *T. gondii*-forming vacuole called the parasitophorous vacuole (PV) inside infected cells and destroy the structure, leading to parasite killing. IFN-γ -inducible GTPases involving anti-*T. gondii* cell-autonomous immunity consist of p47 immunity-related GTPases (IRGs) and p65 guanylate binding proteins (GBPs) (Howard et al., 2011; Yamamoto et al., 2012; Saeij and Frickel, 2017). Most IRGs and GBPs are recruited to PV membranes (PVM) and cooperatively disrupt the membrane structure.

Sequential and hierarchical recruitment of IRGs and GBPs leads to efficient PVM disruption and pathogen clearance (Khaminets et al., 2010). The IRG Irgb6 has been shown to be localized at the PVM soon after *T. gondii* invasion to host cells and acts as a pioneer for the recruitment of other IRGs and GBPs (Khaminets et al., 2010; Lee et al., 2019). Genetic ablation of Irgb6 results in severely impaired accumulation of other IRGs and GBPs, reflecting the pioneering role of Irgb6 to induce host defense (Lee et al., 2019).

Recent reports of crystal structures of Irga6 in various nucleotide states and Irgb10 in the GDP state elucidated the basic architecture of IRGs, consisting of a GTPase domain and N-terminal and C-terminal helical domains (Ghosh et al., 2004; Ha et al., 2021). Structural studies also indicated that homo-dimerization through the GTPase domain interface is required to activate the GTPase of IRG proteins (Pawlowski et al., 2011; Schulte et al., 2016; Ha et al., 2021). However, the structural mechanism of PVM recognition is still vague. Irga6 and Irgb10 utilize a myristoylated glycine at their N-terminus to attach to the PVM (Haldar et al., 2013), although detailed knowledge of the N-terminal structure is missing due its flexibility. Irgb6 does not have the myristoylated glycine, and instead recognizes phospholipids such as phosphoinositide 5P (PI5P) and Phosphatidylserine (PS) via the C-terminal amphipathic α-helices in order to bind present in the PVM (Lee et al., 2019). Due to the lack of structural information on Irgb6, however, the structural basis for such phospholipid recognition and its relationship with nucleotide binding remain unclear.

Here, we aimed to elucidate the PVM recognition mechanism of Irgb6 by X-ray crystallography. We further investigated the membrane binding interface by *in silico* phospholipid docking, followed by validation using mutational analyses.

## Results

### Overall architecture of the Irgb6 monomer in two distinct nucleotide states

To explore the atomic structure of Irgb6, full length mouse Irgb6 was expressed and purified. Size exclusion chromatography (SEC) of purified Irgb6 produced two peaks (Fig. S1 A). SDS-PAGE analysis indicated that Irgb6 was mainly eluted in the second peak. Considering that the estimated molecular weight of Irgb6 is 47.3 kDa, the eluted Irgb6 in the second peak should be a monomer. We also examined the GTPase activity of purified Irgb6 through anion exchange chromatography, indicating retained GTPase activity of Irgb6 (Fig. S1 B).

We successfully crystallized the monomer fraction of Irgb6. As indicated by the SEC analysis, one molecule of Irgb6 was included in the unit cell. The atomic structures of mouse Irgb6 monomer were solved in two states--with GTP and without nucleotide (nucleotide free: NF)--at 1.5 and 1.9 Å resolution, respectively (Fig. 1, A-B and Fig. S1 C). The former structure possesses the GTP and Mg^2+^ ion in the nucleotide binding pocket (GTP-bound Irgb6) (Fig. S2 A). The latter does not have any nucleotide or ion in the pocket (NF Irgb6) (Fig. S2 B).

**Figure 1.**
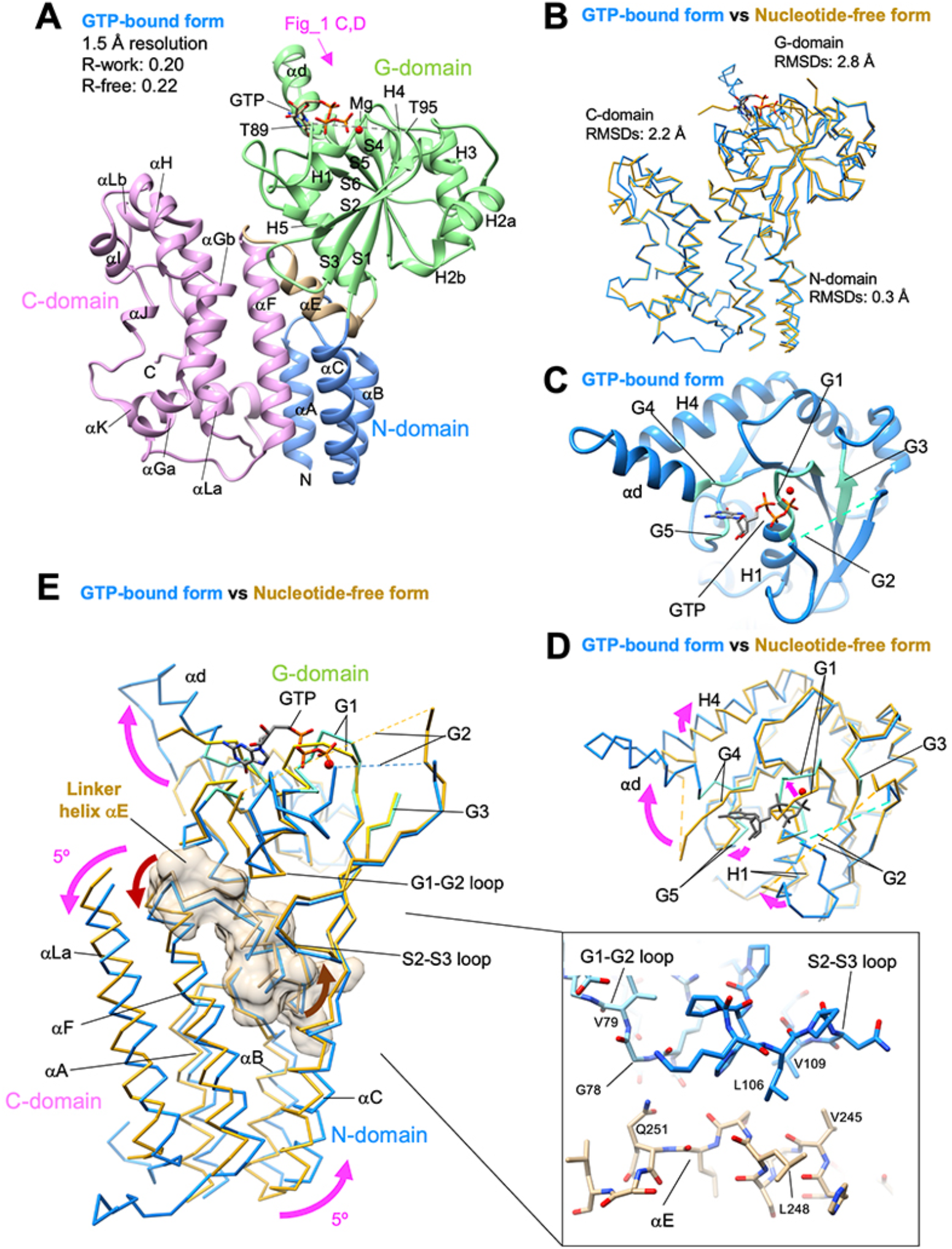
Crystal structures of Irgb6 with GTP and without any nucleotide. (A) Crystal structure of Irgb6 with GTP. (B) Structural comparison between Irgb6 with GTP (blue) and without any nucleotide (yellow-brown), superimposed using all residues to minimize RMSDs. (C) Structure around the nucleotide-binding pocket in G-domain observed from the top indicated by pink arrow in panel (A). (D) Conformational change of nucleotide binding pocket during GTP-binding. (E) Conformational change of N-and C-domains during GTP-binding. Irgb6 with GTP (blue) and without any nucleotide (yellow-brown) was superimposed on their G-domains to illustrate the relay of conformational changes from the nucleotide-binding pocket. Whole N-domain, main components of G-domain around the nucleotide binding pocket, linker helix αE, and helices G and La of C-domain are shown. Linker helix αE is also shown with surface model. (Inset) Close-up view of the interactions between switches I-II and the helix αE.

The overall architectures of Irgb6 are similar with previously solved Irga6 or Irgb10 structures (Ghosh et al., 2004; Ha et al., 2021). They consist of an N-terminal helical domain (N-domain; amino acids 1-55; αA– αC) (blue in Fig. 1 A), a GTPase domain (G-domain; amino acids 56-239; H1–H5, αd, S1–S6) (green in Fig. 1 A), and a C-terminal helical domain (C-domain; amino acids 255-415; αF– αL) (pink in Fig. 1 A) (Fig. S2 C). Helix αE serves as a linker among the three domains (amino acids 240-254) (brown in Fig. 1 A). The G-domain of Irgb6 exhibits a dynamin-like α/β structure with a central β-sheet surrounded by helices on both sides. The N-and C-domains stand side by side and are composed of 11 helices, most of which align parallel or anti-parallel.

### Nucleotide dependent conformational change of Irgb6

The Irgb6 structures in two distinct nucleotide states take on similar conformations with overall root-mean square deviations (RMSDs) of 2.4 Å (Fig. 1 B). The N-domain is apparently in the same conformation, with RMSDs of 0.3 Å, demonstrating no conformational change observed during GTP-binding. On the other hand, the G-and C-domains change their conformation significantly with overall RMSDs, of 2.8 and 2.2 Å, respectively.

The conformational changes of the G-domain are concentrated around the nucleotide binding pocket, consisting of five consensus sequences among p47 GTPases (green in Fig. 1 C and Fig. S2 C). The G1/P-loop (GxxxxGKS) in the GTP-bound state recognizes α-, β-phosphates of GTP and the Mg^2+^ ion, while that in the NF state override the corresponding binding site of the α-, β-phosphates, kicking the GDP out from the nucleotide-binding pocket of Irgb6 (Fig. 1 D and Fig. S2, A-B). The G2/switch I region follows the G1/P-loop and helix H1. Thus, the conformational change of G1/P-loop directly transduces changes in H1 and G2/switch I, although the G2 loop was almost invisible in both structures because G2 did not coordinate to the γ -phosphate or Mg^2+^ ion (Fig. 1, C-D and Fig. S2, A-B). Conformational changes were not apparent in the G3/switch II region (Fig. 1 D). Switches I and II are necessary for the hydrolysis of GTP by coordinating the γ- phosphate or Mg^2+^ ion. This γ- phosphate recognizing reaction is referred to as an “isomerization”. In both Irgb6 structures solved here, however, neither switches I nor II coordinate to GTP or Mg^2+^; thus, our GTP-bound Irgb6 takes on a pre-isomerization state. (Fig. 1, C-D and Fig. S2 A) (Nitta et al., 2004; 2008). The Mg^2+^ ion is thus unstable, represented by its high B-factor value (B-factor: Mg^2+^ 42.9 Å^2^; Mean 36.59 Å^2^). Consistently, stable coordinated waters for Mg^2+^ were not observed even in the 1.5 Å resolution map (Fig. S2 A).

The G4 and G5 regions recognize the base of GTP (Fig. 1 C). The G4 and following helices αd and H4 change their conformation largely from the NF state. The αd of the GTP-bound state forms an α-helix by a loop-to-helix transition from the NF state, inducing a clockwise rotation of helix H4 (Fig. 1 D). It should be noted that, in the Irga6 or Irgb10 structure, these helices αd and H4 serve as an interface for homo-dimerization (Pawlowski et al., 2011; Schulte et al., 2016; Ha et al., 2021). Homo-dimerization is thought to be required to activate the GTPase of IRGs. By analogy with Irga6 and Irgb10, therefore, nucleotide-binding or release appear to initiate or break homo-dimerization of Irgb6, respectively, to control the GTPase activity of Irgb6.

A conformational change in the C-domain was observed around helices αH, αI, and αLb (Fig. 1, A-B). These helices do not contact directly to either the G-domain or the neighboring molecule in the crystal packing environment. Thus, how these conformational changes are induced by nucleotide binding is still not clarified.

Finally, we examined the inter-domain rearrangement from the NF state to the GTP-bound state by superimposing two structures on their G-domains. As a result, N-and C-domains cooperatively rotated 5° in a counterclockwise direction around the G-domain, during GTP-binding (Fig. 1 E). The linker helix αE plays a pivotal role to transduce this nucleotide-dependent conformational change. It makes contacts not only with the preceding loop to switch I (G1-G2 loop) through hydrogen-bonds between the main chains and Gln251, but also with the S2-S3 loop preceding to the switch II region through the hydrophobic residues Leu106, Val109, Val245, and Leu248 (inset of Fig. 1 E). Therefore, the helix αE can sense the conformational change of two switch regions and transduce the change to the N-and C-domains. From the NF state to the GTP-bound state observed here, the conformational change of switch I pushes the αE toward the helical domains, generating rotational change<u>s</u> in the N-and C-domains (Fig. 1 E).

### Comparison of Irgb6 structures with Irga6 and Irgb10 structures

We next compared the newly-solved Irgb6 structures with previously-solved IRG structures. Irga6 structures were reported in three states with GMPPNP (PDB ID: 1TQ2), GDP (PDB ID:1TPZ), and without any nucleotide (PDB ID: 1TQD) (Ghosh et al., 2004). The Irgb10 structure was reported in the GDP state (PDB ID: 7C3K) (Ha et al., 2021). We thus compared our Irgb6 structures with these four structures. Figure 2 A shows RMSDs of N-, G-, C-domains among six structures. This comparison can be summarized in three main findings: 1) The N-domain assumes a very similar structure among IRGs except for the N-terminal end; 2) the structural similarities of the G-domain reflect the nucleotide state of IRGs; 3) the C-domain exhibits large structural variation among IRGs.

**Figure 2.**
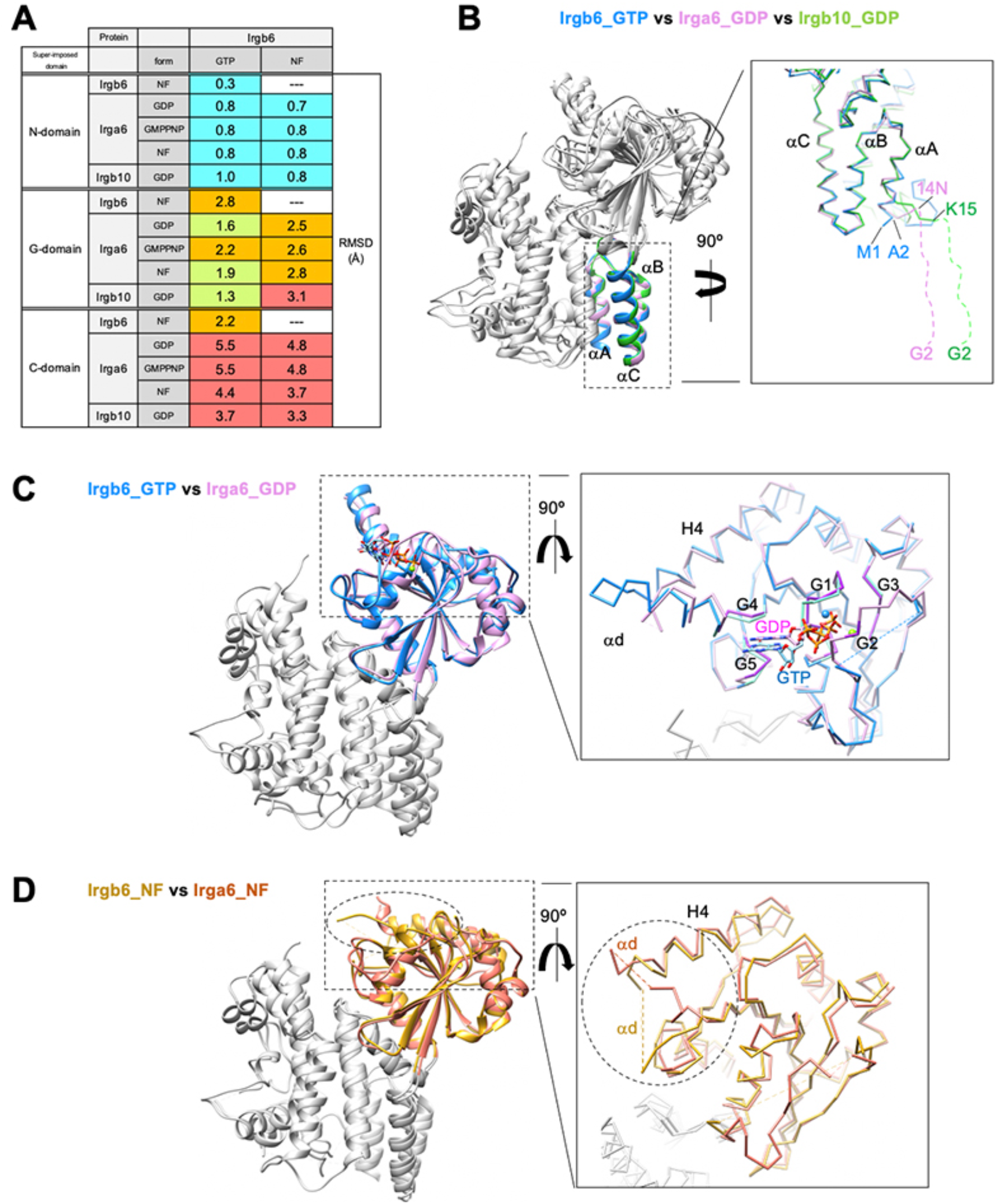
Structural comparison among Irgb6, Irga6, and Irgb10. (A) RMSDs among Irgb6, Irga6, and Irgb10, superimposed on their N-, G-, and C-domains. RMSDs 1: cyan, 1 RMSDs 2: yellow-green, 2 RMSDs 3: orange, 3 RMSDs: red. (B) Structural comparison among Irgb6 with GTP (blue), Irga6 with GDP (pink), and Irgb10 with GDP (green) superimposed on their N-domains. (C) Structural comparison between Irgb6 with GTP (blue) and Irga6 with GDP (pink), superimposed on their G-domains. (D) Structural comparison between Irgb6 without any nucleotide (yellow-brown) and Irga6 without any nucleotide (red), superimposed on their G-domains.

The N-domains of the six structures take on very similar conformations, with RMSDs less than 1.0 Å (Fig. 2, A-B and Fig. S2 D). However, marked difference between Irgb6 and the others exists. The N-terminal helix αA of Irgb6 is slightly longer than those of Irga6 and Irgb10 (Fig. 2 B and S2 D). Instead, Irga6 and Irgb10 have an approximately 15-residue addition before helix αA (Fig. S2 C). This includes the N-terminal glycine residue which is crucial for PVM localization of Irga6 and Irgb10. Gly2 is known to be myristoylated, which allows it to bind to the PV membrane. Since Irgb6 does not have this additional sequence or an equivalent glycine, a different mechanism for recruiting Irgb6 to the PVM exists, as detailed below.

The G-domain of the GTP-bound Irgb6 is most similar with the GDP form of Irga6 (RMSDs = 1.6 Å) or Irgb10 (RMSDs = 1.3 Å), rather than the GMPPNP form of Irga6 (RMSDs = 2.2 Å) (Fig. 2 A and 2 C). This is reasonable because the G1, G4 and G5 sequences recognize α-, β-phosphates and nucleotide base; whereas, γ- phosphate is not trapped by G2/G3 sequences (Fig. 1 C and Fig. S2 A). In our GTP-bound structure, therefore, Irgb6 only recognizes a “GDP part” of GTP so that it resembles the GDP form of IRG proteins. By analogy with Irga6 or Irgb10 (Ghosh et al., 2004; Ha et al., 2021), homo-dimerization of Irbg6 might trigger the isomerization of the G-domain to assume the active GTP form.

The G-domain of NF Irgb6 exhibits a similar conformation to the NF form of Irga6, although the helix H4 and surrounding structures assume different conformations (Fig. 2 D). This difference can be explained by the increased flexibility of G4 due to the absence of nucleotide, which is further stabilized by a neighboring molecule in the crystal packing environment (Fig. S2 E). Therefore, the structure of the G-domain and its conformational changes during the GTPase cycle are basically conserved among IRGs.

## Unique conformation of C-domain in Irgb6 structures

In comparison with the N-and G-domains, the structural similarity of the C-domain is low among IRGs (Fig. 3 A). The RMSDs between different subfamilies of IRGs are greater than 3Å (Fig. 2 A). The C-terminal 22 residues of the C-domain helix αLb and the following tail, are unique additions in Irgb6 (Fig. S2 C). The αLb helix is rich in basic residues, whereas the tail is rich in acidic residues. The biological significance of the C-terminal tail is currently unknown.

**Figure 3.**
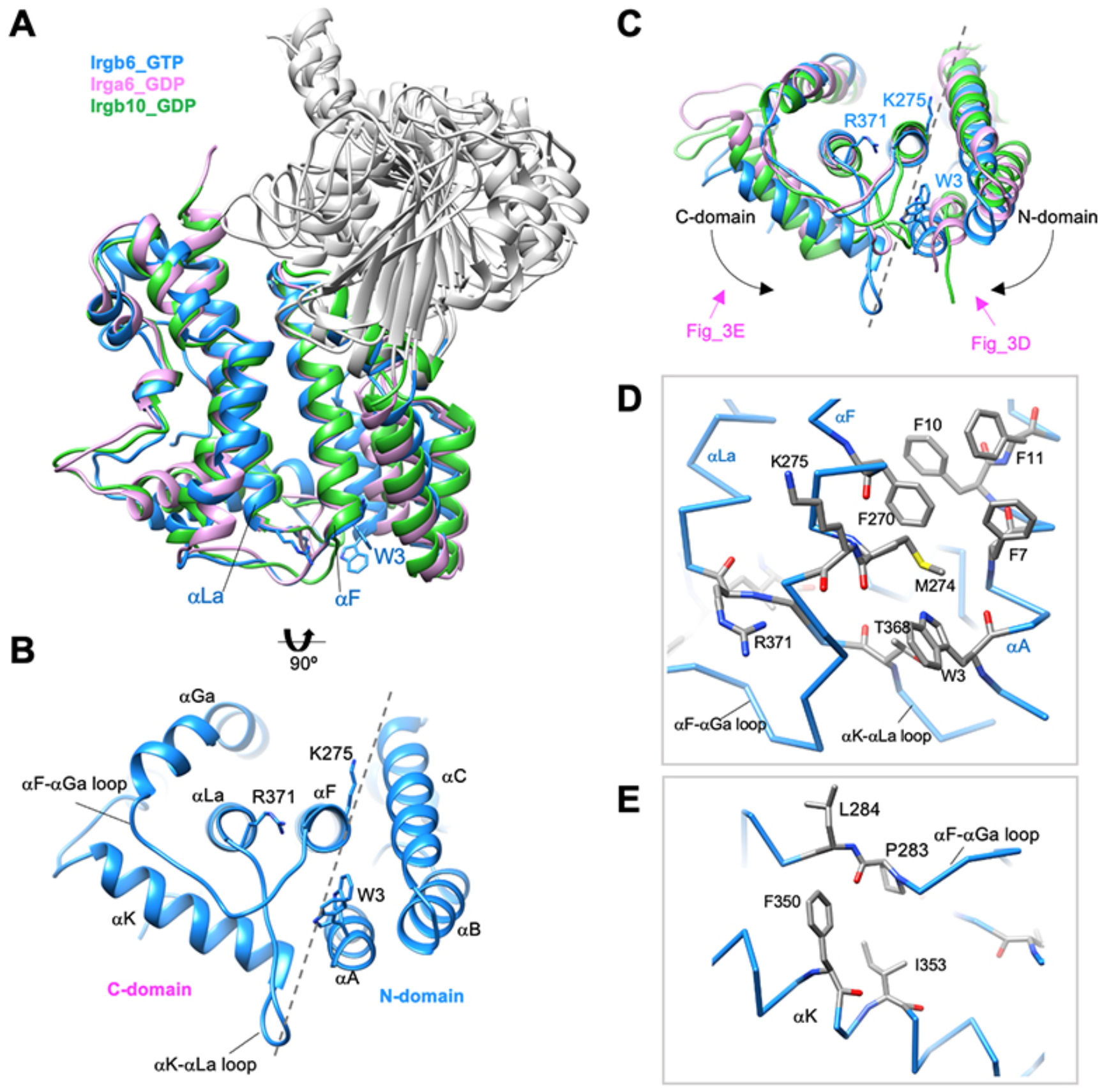
Unique conformation of C-domain in Irgb6. (A) Structural comparison among Irgb6 with GTP (blue), Irga6 with GDP (pink), and Irgb10 with GDP (green) superimposed on their C-domains. (B) Bottom view of Irgb6. Broken line indicates the boundary between N-and C-domains. (C) Bottom view of panel (A) showing the conformational differences among Irgb6 with GTP (blue), Irga6 with GDP (pink), and Irgb10 with GDP (green). (D) Close-up view of the boundary indicates the interaction between the helices αA and F. (E) Close-up view of the interaction between the αF-αGa loop and the helix αK.

There are two antiparallel long helices, αF and αLa, which take on well conserved conformations among IRGs, penetrating the N-and C-domains (Fig. 3 A). Observed from the bottom side, the other side of the GTP-pocket, the helix pair is located at the center of the N-and C-domains, surrounded by five helices from the N-and C-domains (helices αA, αB, αC in N-domain and helices αGa and αk in C-domain) (Fig. 3 B). Two connecting loops, the αF-αGa loop and αK-αLa loop, extend from the central helix pair. These loops, as well as the surrounding five helices, change the conformation significantly among IRGs (Fig. 3 C). The C-domain helices rotate counterclockwise around the central pair, whereas the N-domain helices rotate toward the clockwise direction, thus closing the cleft between N-and C-domains (dashed lines in Fig. 3, B-C). These helices are connected through hydrophobic contacts where residue Trp3 of the αA acts as a keystone (Fig. 3, B and D). Trp3 takes alternative conformations and links two connecting loops with three N-terminal helices, thus contributing to the inter-domain contact between N-and C-domains. Also, the aromatic residues connect the helix αA with αF, supporting the cooperative movement of N-and C-domains (Fig. 3 D). The unique conformation of the αF-αGa loop in Irgb6 is also supported by the hydrophobic residues Phe350 and Ile353 of helix αK, which also assumes a unique conformation because of the long insertion of αK-αLa loop (Fig. 3 E).

Quite suggestively, two basic residues Lys275 and Arg371, which are necessary for PVM recruitment of Irgb6, are located at the ends of the central pair (Fig. 3 B) (Lee et al., 2019). Considering that the myristoylation site of Irga6 and Irgb10 exists at the N-terminal end, close to the end of central pair, these sites were assumed to contribute to the binding of Irgb6 to the PVM.

### Docking simulation of phospholipids to the Irgb6

We previously reported that Irgb6 binds to PI5P and PS, which are both components of the *T. gondii* PVM (Lee et al., 2019). Thus, we simulated the docking of various phospholipids to our Irgb6 structure in order to investigate the specificity. The αF-αGa loop is well defined in GTP-bound Irgb6, so we utilized this structure for the docking experiments.

Molecular docking was performed to investigate the interaction between Irgb6 protein with four phospholipids (PI5P: PubChem 643966, PS: PubChem 9547090, PE: PubChem 160339 and PC: PubChem 445468) using Glide (Halgren et al., 2004). As a consequence, the head groups of phospholipids were docked on the αF-αGa loop and the central helix pair (Fig. 4, A-B). Hereafter, we thus denoted the αF-αGa loop as the “PVM binding loop”. In order to extend sampling of this region, the grid box was approximately centered on residues Trp3, Lys275 and Arg371, with small perturbations, and two rotamer states for Trp3 and Arg371 were independently considered, for a total of six docking runs per ligand. Since Glide measures the ligand-receptor binding free energy in terms of Glide Score, we compared all six Glide Scores of four phospholipids to evaluate their binding affinity to Irgb6. Consistent with our previous report (Lee et al., 2019), the mean Glide scores of the polar head groups indicate that the binding free energy of Irgb6 to the PI5P polar head is lower than that of PS, PE, or PC (Fig. 4 C and S3 A).

**Figure 4.**
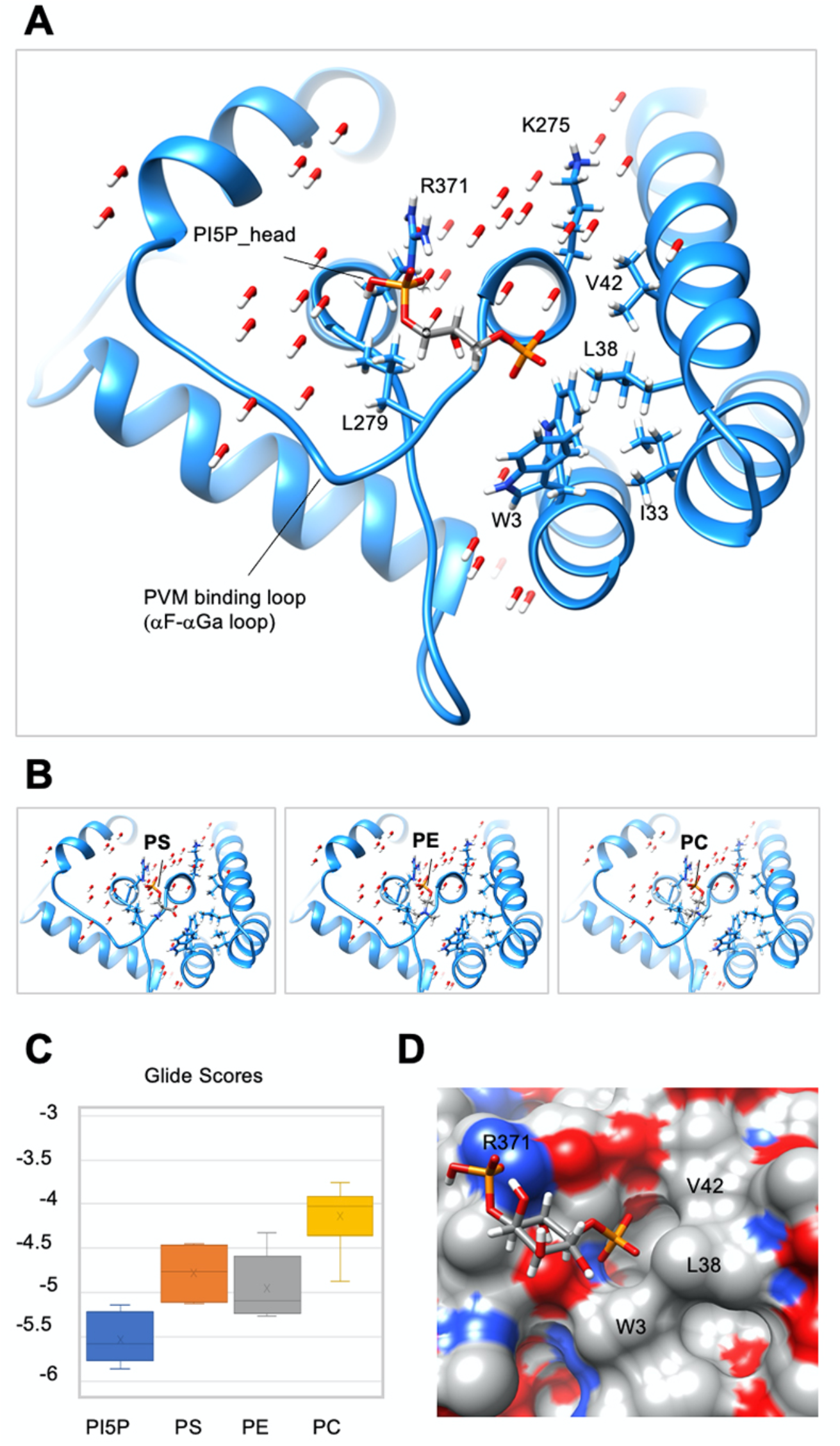
Docking simulation of phospholipids to the Irgb6. (A) Docking of the polar head of PI5P to the GTP-bound Irgb6. (B) Docking of the polar head of PS, PE, and PC to the GTP-bound Irgb6. (C) Glide scores of Irgb6 docking with phospholipid polar head groups. See Figure S5 for detail. (D) Surface presentation of the PI5P pocket colored by elements. Blue, nitrogen, red: oxygen, gray: carbon. The left side of the pocket where the PI5P head docks is covered with the hydrophilic/ionic residues, whereas the right side is covered with the hydrophobic residues (Trp3, Leu38, Val42).

The tips of phosphate groups of PI5P bind directly to Arg371 by hydrogen bonds and form salt bridges and hydrogen bonds to Lys 275 via several water molecules (Fig. 4 A). The Inositol makes hydrophobic contact with Leu279 of the PVM binding loop. The following phosphate faces toward the N-terminal helices, thus the acyl chain would extend toward the N-terminal helices. This surface is mainly covered by hydrophobic residues (Trp3, Ile33, Leu38, Val42), thus environmentally preferable to acyl chain extension (Fig. 4, A and D).

We further checked docking of four phospholipids with several lengths of glycerol backbone or acyl chain to Irgb6. Consequently, Glide Scores of PI5P were always higher than those of PS, PE, or PC, suggesting PI5P is the best suited phospholipid to be targeted by Irgb6 (Fig. S3, B-C). It should be noted that the short glycerol backbone or short acyl chains tended to direct themselves towards the hydrophobic pocket near Trp3. Considering that the lipid tails should be embedded in the bilayer membrane (Muftuoglu et al., 2016) some conformational change at the N-terminal helices should occur before rigid binding of Irgb6 to the PV membrane.

### *In vivo* evaluation of structural model for PVM binding

Docking simulations indicated that the head group of PI5P is on the PVM binding loop and the central helix pair, thus the acyl chain should run towards the N-terminal helix αA. To assess the role of the putative membrane binding region in Irgb6, we generated three Irgb6 mutants. A part or all of the PVM binding loop was substituted with that of Irga6. The Irgb6 mutant in which the 277-286 amino acids were entirely substituted with those of Irga6 was denoted Irgb6_a6(all). The Irgb6 mutant in which a cluster of glycine residues, Gly277, Gly285 and Gly286, were substituted with aspartic acid, threonine, and phenyl alanine, respectively, was denoted Irgb6(G277D/G285T/G286F). In addition, a point mutant, in which the tryptophan at position 3 at the N-terminus of Irgb6 was substituted with alanine (Irgb6 W3A), may affect the structure of the membrane binding pocket.

We reconstituted wildtype Irgb6 and the Irgb6_a6(all), the G277D/G285T/G286F, or the W3A mutant mutants in Irgb6-deficient MEFs (Fig. 5 A). We confirmed that wild-type and mutant Irgb6 proteins were expressed at comparable levels in the reconstituted cells (Fig. 5 A). Then we tested them for IFN-γ- induced reduction of *T. gondii* numbers and the recruitment to *T. gondii* PVM (Fig. 5, B-D). When IFN-γ- induced killing activity was examined, Irgb6-deficient MEFs reconstituted with wild-type Irgb6 were able to recover the killing activity (Fig. 5 B). In sharp contrast, Irgb6 KO MEFs that expressed the Irgb6_a6(all), the G277D/G285T/G286F, or the W3A mutants were not able to restore this killing activity (Fig. 5 B). Furthermore, reconstitution of wildtype Irgb6 in Irgb6-deficient MEFs recovered the recruitment to *T. gondii* PVM, whereas that of the Irgb6_a6(all), the G277D/G285T/G286F, or the W3A mutants did not reconstitute the mutant recruitment (Fig. 5 D). Collectively, a cluster of glycine residues in the PVM biding loop and the N-terminal tryptophan are essential for the Irgb6 PVM targeting and its killing function.

**Figure 5.**
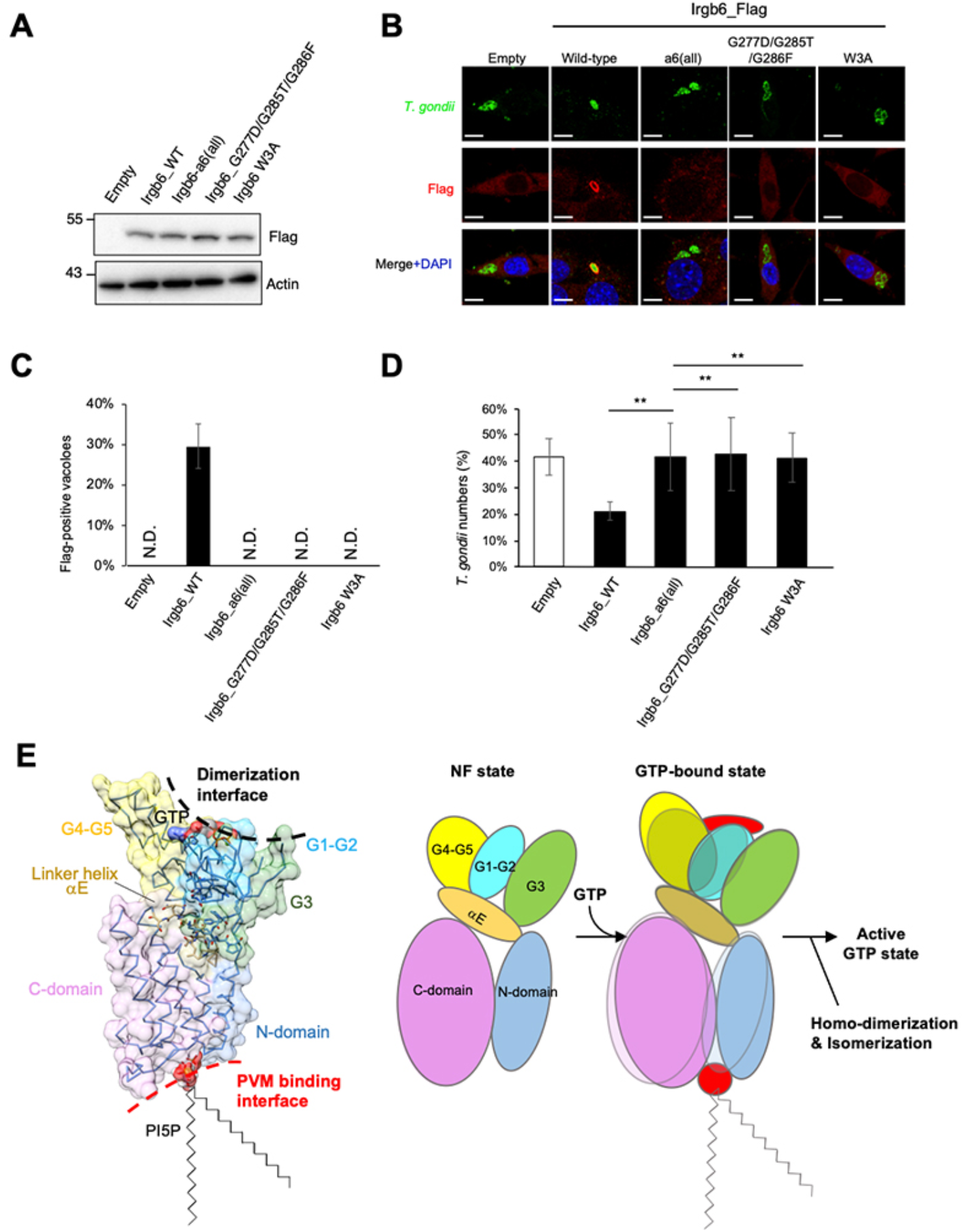
The membrane binding region is essential for Irgb6 accumulation on *T. gondii* PVM. (A) Western blot image to detect stably expressed Irgb6 protein after retroviral transfection and puromycin selection. (B) Confocal microscope images to show the localization of Irgb6-Flag (red) to *T. gondii* PV (green), and DAPI (blue) at 4 h post infection in IFN-γ treated Irgb6-KO MEFs reconstituted with indicated Irgb6. (C) Recruitment percentages of Irgb6_Flag. (D) *T. gondii* survival rate in the indicated Irgb6 reconstitution in Irgb6 KO MEFs with IFN-γ stimulation relative to those without IFN-γ treatment by luciferase analysis at 24 h post infection. All graphs show the mean ± SEM in three independent experiments. All images are representative of three independent experiments. N.D., not detected; \*\**P* < 0.01. T. gondii survival and Irgb6_Flag recruitment comparison between genotypes applied one-way ANOVA (Tukey’s multiple comparisons test). White arrows to indicate recruitment of effector on *T. gondii* PV. Scale bars on microscope images represent 10 μm. (E) Structural model of PVM recognition during the GTP-binding.

## Discussion

Irgb6 has a crucial role to target the PVM of *T. gondii* and destroy them. Our atomic structures of Irgb6 solved here elucidated the structural mechanisms of PVM recognition by Irgb6. Irga6 or Irgb10 was reported to have the myristoylation site at their N-terminus (Haldar et al., 2013). Irgb6 utilizes different mechanisms in which Irgb6 binds to PI5P or PS to assess the PVM. The PI5P binding site is located at the bottom surface that is composed of both the N-and C-domains, opposite to the GTP-binding pocket (Fig. 5 E).

Irgb6 maintains structural features common among IRGs. It is composed of three domains, N, G, and C; and, the nucleotide dependent conformational change is also conserved in comparison to Irga6 structures. Our structures solved here do not represent the active GTP form in the pre-hydrolysis state. However, the purified Irgb6 protein used for the crystallization had GTPase activity (Fig. S1 B), suggesting that Irgb6 without any accessory proteins or co-factors can hydrolyze GTP. Thus, by analogy with Irga6 (Pawlowski et al., 2011), homo-dimerization of Irgb6 using the G-domain as an interface will activate the Irgb6 GTPase.

Nucleotide dependent conformational change of G-domain transduces N-and C-domains through the linker helix αE (Fig. 5E). This helix is located at the center among N-, G-, and C-domains and relays the conformational change of G2/switch I and G3/switch II to the N-and C-domains to generate the rotational movement known as a “power stroke” (Chappie et al., 2011) Our structures solved here represents the large conformational change of G4-G5 during GTP-binding, making the top interface ready for homo-dimerization (Fig. 5 E). They also show small rotation of N-and C-domains via the helix αE movement that is induced by the small change of G2/switch I (Fig. 5 E). By the analogy with Irga6, homo-dimerization through the G-domains would change and stabilize the conformation of switch I and switch II. It would induce a power stroke of N-and C-domains relayed through helix αE, thus remodeling the PV membrane. Further structural studies are needed to prove this hypothesis.

In contrast to the highly conserved GTPase domain, the helix αA in the N-domain and whole C-domain, both of which serve as a membrane binding interface through PI5P binding, have large conformational variations among IRGs. In comparison with Irga6, the N-and C-domains of Irgb6 rotate in opposite directions, resulting in closure of the cleft between N-and C-domains (Fig. 3, B-C). The class specific tryptophan 3 residue at the N-terminus plays a pivotal role for the cleft closure between N-and C-domains, mainly made by the hydrophobic interactions. Due to this movement, Irgb6 can produce an Irgb6-specific membrane-binding interface to recognize the PVM.

As described above, if we used the head domain of PI5P with the short acyl chain for the docking simulation, the acyl chains tended to direct themselves towards the hydrophobic pocket between the N-and C-domains. The acyl chain of the membrane-bound PI5P, on the other hand, must be oriented toward the PVM. To understand the conformational change during GTP binding through homodimerization, we compared the membrane binding surface of Irgb6 with that Irga6 in the active GTP form in which conformational rearrangements of N-and C-domains were exclusively observed in Irga6. Interestingly, Irga6 had a wider open pocket than that of Irgb6 precisely at the putative acyl chain binding side of PI5P (Fig. S3 D). In this conformation, the acyl chain could interact hydrophobically with Irga6 and then pass through the pocket, and then enter the membrane. We further checked the hydrophobic pocket of Irgb6. GTP-bound Irgb6 takes an alternative conformation at Trp3. In these two alternative forms, the depths or sizes of the pocket openings differ significantly (Fig. S3, E-F). Thus, the conformation of the N-terminal helix could alter the shape and depth of hydrophobic pocket of Irgb6. From these observations, we assume that the GTPase activation by homodimerization of Irgb6 could change the helical domain to open the hydrophobic pocket in order to accommodate the acyl chain. Further structural studies are required to prove this hypothesis.

In the present study, we solved here the atomic structure of Irgb6 monomer and elucidated the structural detail of PVM binding interface of Irgb6. Considering that Gbp1 regulates the localization or activity of Irgb6 on the PVM, the biochemical and structural analyses of Gbp1–Irgb6 interaction are required to solve the molecular mechanisms of PVM disruption and pathogen clearance (Khaminets et al., 2010). Also, the rhoptry protein 18 (ROP18), a serine threonine kinase secreted by *T. gondii*, phosphorylates threonine residues in switch I of Irgb6 to disarm the innate clearance by host cells (Fentress et al., 2010). By elucidating the structural mechanisms of how ROP18 inactivates Irgb6, therefore, the whole picture of host cell-autonomous immunity and microbial counter-defense system will be unveiled.

## Materials and Methods

### Protein expression

The full-length *Mus musculus* Irgb6 gene (*Tptg 2*, Gene ID: 100039796) was PCR-amplified with specific primers (5′-<u>GAAGTTCTGTTCCAGGGGCCC</u>ATGGCTTGGGCCTCCAGC -3′ and 5′-

<u>CGATGCGGCCGCTCGAG</u>TTATCAAGCTTCCCAGTACTCGG -3′; the original sequence of pGEX-6P-1 are underlined) from the pWT_Irgb6_full (Lee et al., 2019) and then subcloned into the directly downstream of PreScission protease site pGEX-6P-1 (Cytiva) by Gibson Assembly system (*New England Biolabs* Inc.) to create pRN108. The pRN108 was transformed into *E. coli* strain BL21(DE3). The transformant were grown in LB medium with 50 mg/L ampicillin at 25°C to an OD600 nm of 0.4, and GST-tagged Irgb6 was expressed overnight with final 0.1 mM Isopropyl β-D-1-thiogalactopyranoside. The cells were harvested and stored at -80°C.

### Protein Purification

Irgb6 was purified at 4°C. The frozen BL21(DE3) cells were suspended in solution-I (50 mM HEPES-KOH, pH 7.5, 150 mM NaCl, 2 mM dithiothreitol, 0.7 μM leupeptin, 2 μM pepstatin A, 1 mM phenylmethylsulfonyl fluoride and 2 mM benzamidine) and sonicated on ice. The cell lysate was centrifuged (80,000 g, 30 min) and GST-Irgb6 in the soluble fraction was purified by affinity chromatography using a Glutathione Sepharose 4B column (Cytiva) equilibrated with solution-I. The GST domain of the protein was cleaved by overnight incubation with GST-tagged HRV 3C protease (homemade) on the resin. The free Irgb6 which contains two extra N-terminal residues, Gly-Pro, was eluted with solution-I and was concentrated to with an Amicon Ultra 10-kDa MWCO concentrator (Merck Millipore). The protein was further subjected to size exclusion chromatography on a HiLoad 16/600 Superdex 75 column pg column (Cytiva) equilibrated in solution-II (50 mM HEPES-KOH, pH 7.5, 150 mM NaCl, 5 mM MgCl_2_, 2 mM dithiothreitol). Peak fraction containing Irgb6 at ∼47 kDa elution position was concentrated using the concentrator for crystallization. Protein concentration was estimated by assuming an A280 nm of 0.916 for a 1 mg/ml solution.

### Crystallization

Nucleotide-free Irgb6 crystals diffracting to 1.9 Å resolution were obtained from sitting drops with a 12 mg/ml protein solution and a reservoir solution consisting of 0.1 M MIB buffer pH 6.0 (Molecular Dimensions), 25% Polyethylene Glycol 1500 (Molecular Dimensions) at 20°C. GTP binding Irgb6 crystals diffracting to 1.5 Å resolution were obtained from sitting drops with a 9 mg/ml protein solution containing 2 mM GTP (Roche) and a reservoir solution consisting of 0.1 M Sodium Citrate buffer pH 5.4 (Wako), 18% (w/v) Polyethylene Glycol 3350 (Sigma) at 20°C. GMPPNP binding Irgb6 crystals diffracting to 1.6 Å resolution were obtained from sitting drops with a 9 mg/ml protein solution containing 2 mM GMPPNP (Jena Bioscience) and a reservoir solution consisting of 0.1 M Sodium citrate buffer pH 5.6, 22% (w/v) Polyethylene Glycol 3350 at 20°C.

### Data collection and Structure determination

Single crystals were mounted in LithoLoops (Protein Wave) with the mother liquor containing 10% (v/v) or 20% (v/v) glycerol as a cryoprotectant and were frozen directly in liquid nitrogen prior to X-ray experiments. Diffraction data collection was performed on the BL32XU beamline at SPring-8 (Hyogo, Japan) using the automatic data collection system ZOO (Hirata et al., 2019). The diffraction data were processed and scaled using the automatic data processing pipeline KAMO (Yamashita et al., 2018). The structure was determined using PHENIX software suite (Liebschner et al., 2019). Initial phase was solved by molecular replacement using PDB ID: 1TQD, 1TQ2 and 1TPZ with phenix.phaser. The initial model was automatically constructed with phenix.AutoBuild. The model was manually built with Coot (Emsley and Cowtan, 2004) and refined with phenix.refine. The statistics of the data collection and the structure refinement are summarized in Table S1. UCSF Chimera (Pettersen et al., 2004) was used to create images, compare structures, and calculate Root Mean Square Deviations.

### Analysis of nucleotide component

Irgb6 and GTP were prepared 40 mM in 50 mM HEPES-KOH, pH 7.5, 1 mM MgCl_2_, 1 mM EGTA-KOH, pH7.0, 150 mM NaCl. A 25 μl Irgb6 sample were mixed to equal volume of GTP sample and incubated at 36°C for 30 minutes. A 1 ml of 8 M urea was added to the mixture and heated at 95°C for 1 min, followed by ultrafiltration using Amicon Ultra-0.5 10-kDa MWCO concentrator (Merck Millipore). A 900 μl of the solution that passed through the ultrafiltration membrane was analyzed by anion exchange chromatography using a Mono Q 5/50 GL column (Cytiva) equilibrated with 50 mM HEPES-KOH, pH 7.0. Components of the reaction mixture, GTP and GDP, were completely separated by elution with 0–0.2 M NaCl gradient in 50 mM HEPES-KOH, pH 7.0. Fresh GTP (Nacalai Tesque) and GDP (WAKO) were used to confirm the elution position. A control experiment was performed using the reaction buffer.

### *In situ* Docking simulation

Molecular docking was performed using Schrödinger suite. The 2D structures of the 4 phospholipid ligands, PI5P, PS, PE and PC, were obtained from PubChem (https://pubchem.ncbi.nlm.nih.gov/) (Fig. S4). Acyl chains were truncated up to their corresponding polar head groups. Ligands were also prepared with the polar heads and glycerol backbones, as well as with 4 and 16 carbon acyl chains. The free ligands were converted to three-dimensional structures and their geometries were optimized with the correct chirality using Ligprep. LigPrep was also used to produce different conformations for each ligand structure. Before docking, the Irgb6 protein was prepared using the protein preparation wizard. Subsequently, a grid box was centered on acids Trp3, Lys275 and Arg371. Three similar grid centers and two positions of Trp3 and Arg371 were independently considered (Fig. S5). The prepared ligands were docked with the preprocessed Irgb6 protein grids using Glide standard precision (SP) docking mode with flexible ligand sampling.

### Cells, Mice, and Parasites

MEFs that lack Irgb6 are described previously (Lee et al. 2019). Irgb6-deficient MEFs were maintained in DMEM (Nacalai Tesque) supplemented with 10% heat-inactivated fetal bovine serum (FBS) (Gibco, Life Technologies), 100 U/ml penicillin (Nacalai Tesque), and 100 μg/ml streptomycin (Nacalai Tesque). The complete medium comprised 10% heat-inactivated FBS in RPMI 1640 medium (Nacalai Tesque). *T. gondii* were parental PruDHX, luciferase-expressing PruDHX. They were maintained in Vero cells by passaging every 3 days in RPMI 1640 supplemented with 2% heat-inactivated FBS, 100 U/ml penicillin, and 100 μg/ml streptomycin.

### Reagents

Antibodies against FLAG M2 (F3165), and b-actin (A1978) were obtained from Sigma-Aldrich.

### Cloning and Recombinant Expression

The region of interest of the cDNA corresponding to the wild-type, indicated point mutants or deletion mutants of Irgb6 (GenBank accession no. NM_001145164) were synthesized from the mRNA of the spleen of C57BL6 mice using primers Irgb6_F 5’ -gaattcaccATGGCTTGGGCCTCCAGCTTTGATGCATTCT-3’ and Irgb6_R 5’ -gcggccgcTCActcgagAGCTTCCCAGTACTCGGGGGGCTCAGATAT-3’.

Irgb6_a6(all), G277D/G285/G286F, and W3A mutants were generated using primers a6(all)_F 5’ -TCTTCCTAGAAGCCATGAAGGCTgacctagtgaatatcatcccttctctgacctttATGATCAGTGATATCTTAGAGAAT-3’ and a6(all)_R 5’ -ATTCTCTAAGATATCACTGATCATaaaggtcagagaagggatgatattcactaggtcAGCCTTCATGGCTTCTAGGAAGA-3’ ; G277D/G285/G286F _F 5’ -GTCTTCCTAGAAGCCATGAAGGCTGacGCATTAGCCACCATTCCACTTaactttATGATCAGTGATATCTTAGAGAATCT-3’ and G277D/G285/G286F _R 5’ -AGATTCTCTAAGATATCACTGATCATaaagttAAGTGGAATGGTGGCTAATGCgtCAGCCTTCATGGCTTCTAGGAAGAC-3’ ; W3A _F 5’ -gaattcaccATGGCTgcGGCCTCCAGCTTTGATGCATTCTTTAAGAATTT-3’ products were ligated into the EcoRI/XhoI site of the retroviral pMRX-Flag expression vector for retroviral infection. The sequences of all constructs were confirmed by DNA sequencing.

### Western Blotting

MEFs were stimulated with IFN-γ (10 ng/ml) overnight. The cells were washed with PBS and then lysed with 1× TNE buffer (20 mM Tris-HCl, 150 mM NaCl, 1 mM EDTA, and 1% NP-40) or Onyx buffer (20 mM Tris-HCl, 135 mM NaCl, 1% Triton-X, and 10% glycerol) for immunoprecipitation, which contained a protease inhibitor cocktail (Nacalai Tesque) and sonicated for 30 seconds. The supernatant was collected, incubated with the relevant antibodies overnight, and then pulled gown with Protein G Sepharose (GE) for immunoprecipitation. Samples and/or total protein was loaded and separated in 10% or 15% SDS-PAGE gels. After the appropriate length was reached, the proteins in the gel were transferred to a polyvinyl difluoride membrane. The membranes were blocked with 5% dry skim milk (BD Difco™ Skim milk) in PBS/Tween 20 (0.2%) at room temperature. The membranes were probed overnight at 4°C with the indicated primary antibodies. After washing with PBS/Tween, the membranes were probed with HRP-conjugated secondary antibodies for 1 hour at room temperature and then visualized by Luminata Forte Western HRP substrate (Millipore).

### Measurement of *T. gondii* numbers by a luciferase assay

To measure the number of *T. gondii*, cells were untreated or treated with IFN-γ (10 ng/ml) for 24 hours. Following the stimulation, the cells were infected with luciferase-expressing PruΔHX *T. gondii* (MOI of 0.5) for 24 hours. The infected cells were collected and lysed with 100 μl of 1× passive lysis buffer (Promega). The samples were sonicated for 30 seconds before centrifugation and 5 μl of the supernatants were collected for luciferase expression reading by the dual-luciferase reporter assay system (Promega) using a GLOMAX 20/20 luminometer (Promega). The *in vitro* data are presented as the percentage of *T. gondii* survival in IFN-γ- stimulated cells relative to unstimulated cells (control).

### Immunofluorescence microscopy

MEFs were infected with *T. gondii* (MOI 5 or 2) after stimulation with IFN-γ (10 ng/ml) for 24 hours. The cells were infected for the indicated time in the respective figures and then fixed for 10 minutes in PBS containing 3.7% formaldehyde. Cells were then permeabilized with PBS containing 0.002% digitonin (Nacalai Tesque) and blocked with 8% FBS in PBS for 1 hour at room temperature. Next, the cells were incubated with antibodies relevant to the experiments for 1 hour at 37°C. After gently washing the samples in PBS, the samples were incubated with Alexa 488-and 594-conjugated secondary antibodies as well as DAPI for 1 hour at 37°C in the dark. The samples were then mounted onto glass slides with PermaFluor (Thermo Fischer Scientific) and observed under a confocal laser microscope (FV1200 IX-83, Olympus). Images are shown at ×1000 magnification (scale bar 10 μm). To measure recruitment rates, 100 vacuoles were observed and the numbers of vacuoles coated with effectors were calculated. The counting was repeated three times (three technical replicates). The mean of the three technical replicates was calculated and shown in each circle. The independent experiments were repeated three times (three biological replicates).

### Statistical analysis

Three points in all graphs represent three means derived from three independent experiments (three biological replicates). All statistical analyses were performed using Prism 9 (GraphPad). In *T. gondii* survival and Irgb6_Flag recruitment assays, ordinary one-way ANOVA was used when there were more than two groups.

### Online supplemental material

Fig. S1 A and B shows that Irgb6 expressed in *E. coli* was purified as a monomer and has GTPase activity. Fig. S1 C shows the summary of the data collection and refinement statistics. Fig. S2 A and S2 B shows the GTP pockets of Irgb6_GTP and Irgb6_NF. Fig. S2 C shows amino acid sequence alignment among Irgb6, Irga6 and Irgb10. Fig. S2 D shows the structural homology between Irgb6_NF and Irga6_NF. Fig S2 E shows the crystal packing of Irgb6_NF. Fig S3 A, B and C shows the Glide scores of Irgb6 protein docking with phospholipids’ polar head, with phospholipids’ polar head and glycerol backbone and with phospholipids that have 4C and 16C acyl chains, respectively. Fig S3 D, E and F shows the difference of hydrophobic pockets between Irgb6 and Irga6.

## Author contributions

Y.S.H., M.Y., D.M.S., R.N. conceived the project. Y.S.H., N.S. Y.S. and R.N. performed structural analysis. A.A.S. and D.M.S. performed docking simulation. A.P., M.S., and M.Y. performed cell-based analysis. All authors discussed the results, and Y.S.H., M.Y., D.M.S., R.N. wrote the manuscript.

## Acknowledgments

We thank K. Chin for assistance and other colleagues for discussions. This work was supported by the Japan Agency for Medical Research and Development (AMED) (JP20fk0108137 (M.Y.), JP20wm0325010 (M.Y.), JP20jm0210067 (M.Y.), JP20am0101108 (D.M.S.), JP20gm0810013 (R.N.)). We acknowledge support from the Japan Society for the Promotion of Science (21K06988 (Y.S.H.), 20B304 (M.Y.), 19H04809 (M.Y.), 19H00970 (M.Y.), 19H03396 (R.N.)). This research was also supported by Platform Project for Supporting Drug Discovery and Life Science Research (Basis for Supporting Innovative Drug Discovery and Life Science Research (BINDS)) from AMED under Grant Number JP19am0101070 (*support number 2057*). We also acknowledge support from the Hyogo Science and Technology Association to R. N.

## Online supplemental material

**Figure S1.**
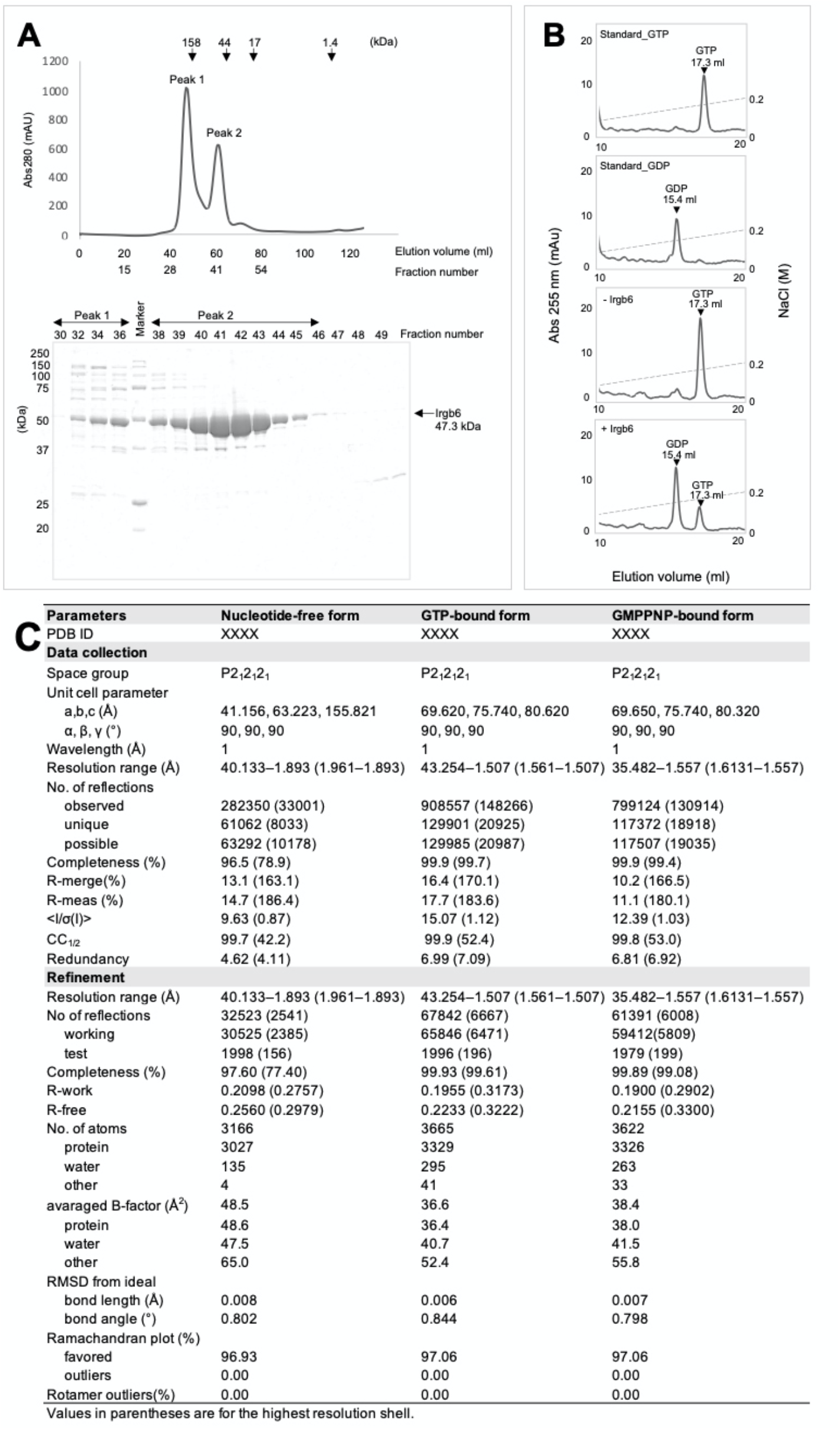
Purification of Irgb6 and Data Collection and Refinement Statistics. (A) SEC analysis of Irgb6. Chromatogram of SEC analysis (top) and SDS-PAGE analysis of peak fractions are shown. (B) Analyses of nucleotide components through the anion column. The incubation of Irgb6 with GTP at 36 °C for 30 min induced hydrolysis of GTP into GDP. (C) Data Collection and Refinement Statistics

**Figure S2.**
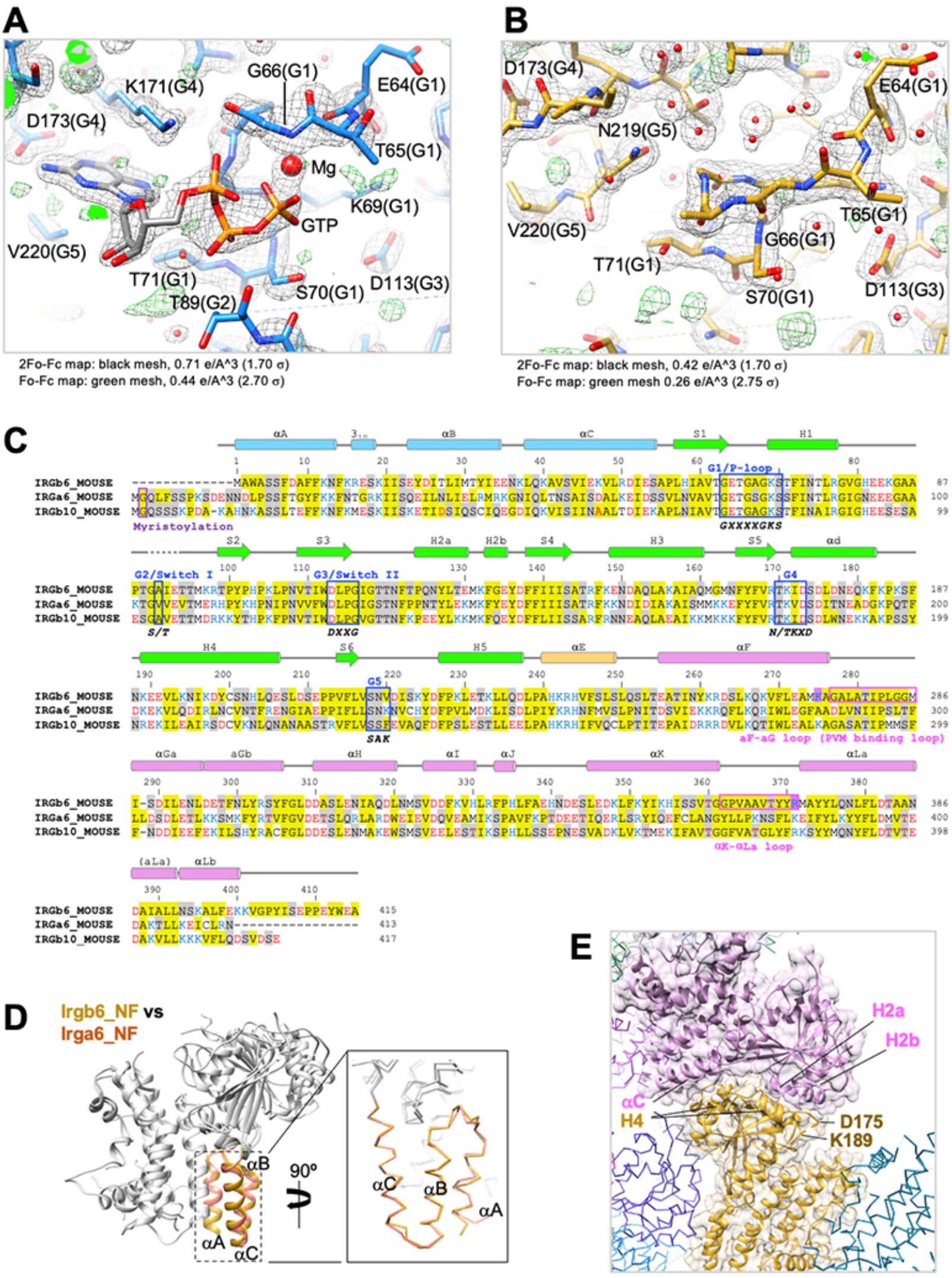
Detailed structures of nucleotide binding pocket, sequence alignment and structural comparison among IRGs. (A) Nucleotide binding pocket of Irgb6 with GTP. (B) Nucleotide binding pocket of Irgb6 without any nucleotide. (C) Amino acid sequence alignment of three mouse IRGs, Irgb6, Irga6, and Irgb10. (D) Structural comparison of N-domains of Irgb6 and Irga6, both in the NF state. (E) Crystal packing environment of Irgb6 in the NF state. The helix H4 was stabilized by the interactions with the neighboring molecule.

**Figure S3.**
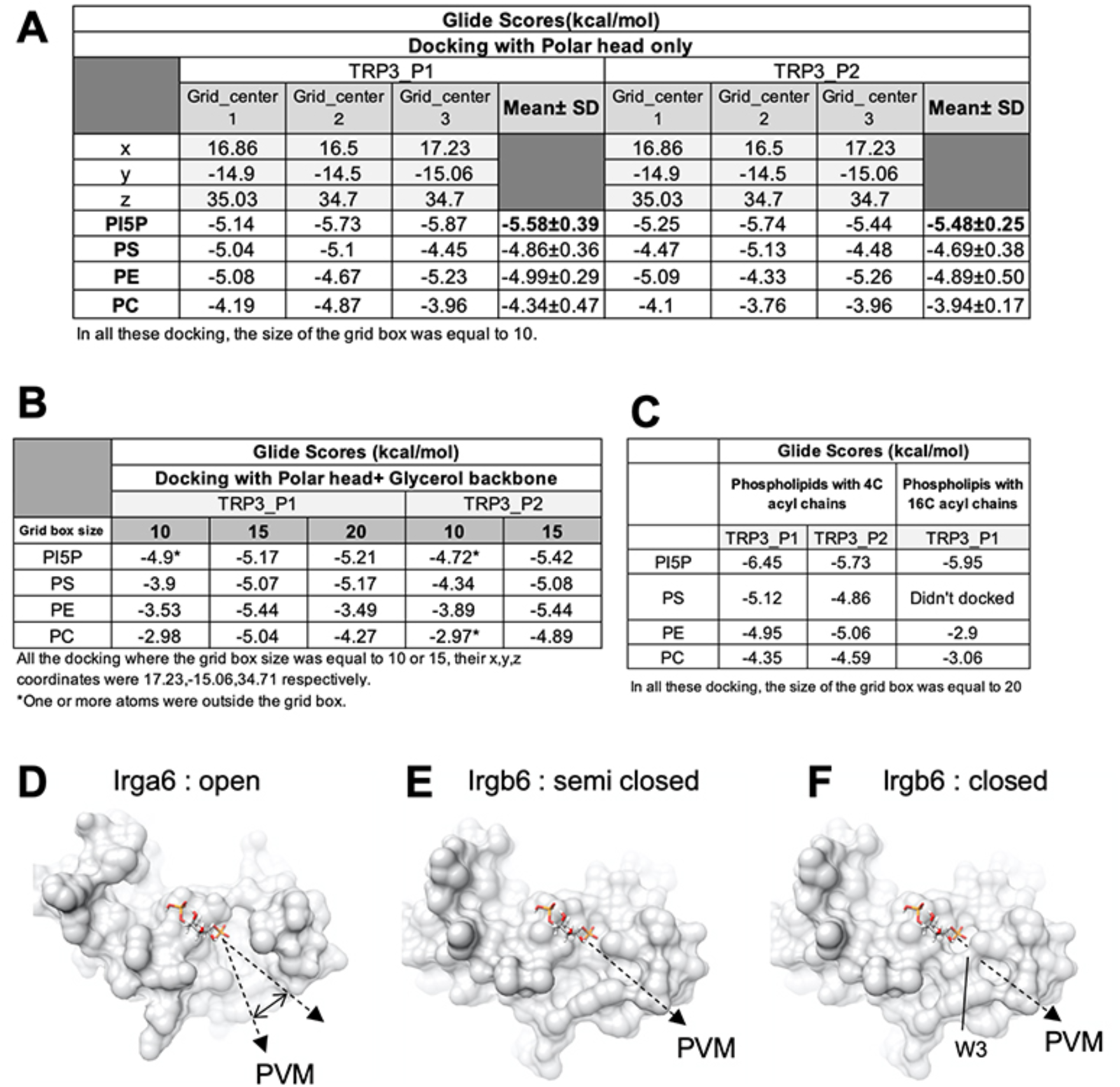
Glide scores of Irgb6 with phospholipids and Structures of PVM-binding pocket. (A) Glide scores of IRGB6 protein docking with phospholipids’ polar head. (B) Glide scores of IRGB6 protein docking with phospholipids’ polar head and glycerol backbone. (C) Glide scores of IRGb6 protein docking with phospholipids that have 4C and 16C acyl chains. (D) PVM-binding site of Irga6 in the active GTP-form (PDB ID: 1TPZ) represents widely open pocket. (E) PVM-binding pocket of Irgb6 in the GTP-bound form (alternative conformation 1) represents semi closed form. (F) PVM-binding pocket of Irgb6 in the GTP-bound form (alternative conformation 2) represents closed form.

## Notes

### Competing Interest Statement

The authors have declared no competing interest.

